# Strain-level diversity impacts cheese rind microbiome assembly and function

**DOI:** 10.1101/652768

**Authors:** Brittany A. Niccum, Erik K. Kastman, Nicole Kfoury, Albert Robbat, Benjamin E. Wolfe

## Abstract

Taxa that are consistently found across microbial communities are often considered members of a core microbiome. One common assumption is that taxonomically identical core microbiomes will have similar dynamics and functions across communities. However, strain-level genomic and phenotypic variation of core taxa could lead to differences in how core microbiomes assemble and function. Using cheese rinds, we tested whether taxonomically identical core microbiomes isolated from distinct locations have similar assembly dynamics and functional outputs. We first isolated the same three bacterial species (*Staphylococcus equorum, Brevibacterium auranticum*, and *Brachybacterium alimentarium*) from nine cheeses produced in different regions of the United States and Europe. Comparative genomics identified distinct phylogenetic clusters and significant variation in genome content across the nine core microbiomes. When we assembled each core microbiome with initially identical compositions, community structure diverged over time resulting in communities with different dominant taxa. The core microbiomes had variable responses to abiotic (high salt) and biotic (the fungus *Penicillium*) perturbations, with some communities showing no response and others substantially shifting in composition. Functional differences were also observed across the nine core communities, with considerable variation in pigment production (light yellow to orange) and composition of volatile organic compound profiles emitted from the rinds (nutty to sulfury). Our work demonstrates that core microbiomes isolated from independent communities may not function in the same manner due to strain-level variation of core taxa. Strain-level diversity across core cheese rind microbiomes may contribute to variability in the aesthetics and quality of surface-ripened cheeses.

## INTRODUCTION

Metagenomic surveys of microbial communities often describe the existence of core microbiomes. Although many definitions currently exist (1), core microbiomes are generally considered to be the set of microbial taxa that are commonly found across all (or many) sampled microbial communities. Many microbiomes, from plant roots to wastewater treatment plants, contain a set of core taxa that are common, highly abundant, and functionally significant (1–4). These core microbiomes can range from just a few species to tens or hundreds of species. For example, most human skin microbiomes are dominated by very similar *Corynebacterium*, *Propionibacterium*, and *Staphylococcus* species (3, 5, 6).

One largely untested assumption is that taxonomically identical core microbiomes will have similar community assembly patterns and functions. More specifically, when comparing 16S rRNA gene sequencing surveys, samples that have very similar compositions of 16S sequences are often assumed to have similar functional potentials. This assumption underlies the development of taxonomy-based microbiome diagnostics and tools used to predict function from taxonomic sequences (7, 8). But independent evolution or coevolution of microbial species within communities may generate previously underappreciated functional diversity across core microbiomes. It is widely accepted that microbial genomes are highly variable within species due to rapid rates of evolution and potential for lateral gene transfer (9–11). Moreover, we know from decades of work in microbial ecology, physiology, and genomics that there is considerable within species trait variation in microbes (12–14). For example, a set of 11 strains of *Brevundimonas alba* isolated from the same freshwater habitat had identical 16S rRNA sequences, but highly divergent carbon utilization profiles and growth rates (15). This intraspecific trait diversity could be ecologically significant, but the impact of strain-level diversity on core microbiome assembly and function is poorly understood (16).

Cheese rinds provide an ideal opportunity to test whether taxonomically identical core microbiomes have similar assembly dynamics and functions and more generally the causes and consequences of core microbiome diversification. Rinds form on the surfaces of cheeses aged in an aerobic environment and are composed of bacteria, yeasts, and filamentous fungi (17–19). Our previous work used amplicon and shotgun metagenomics to describe the bacterial and fungal diversity of 137 cheese rinds from the United States and Europe (17). Three bacterial genera – *Staphylococcus*, *Brevibacterium*, and *Brachybacterium* – were the most frequently detected across cheese rinds and can be considered a core microbiome. Through variation in abiotic and biotic selection pressures applied during cheese production and aging, including abiotic (salinity, pH, resource availability) and biotic (presence of bacterial and fungal neighbors), core cheese microbiomes have the potential to evolve new genotypes and phenotypes with divergent functions.

Here we characterize core microbiome members across cheese rind communities and determine the consequences of core microbiome diversification for community assembly and function. We isolated the same three species of bacteria – *Staphylococcus equorum* (hereafter *Staphylococcus*)*, Brevibacterium auranticum* (hereafter *Brevibacterium*), and *Brachybacterium alimentarium* (hereafter *Brachybacterium*) *-* from nine different cheeses made across the United States and Europe (**Fig. 1A-B**). These nine sets of three co-isolated bacterial species are referred to as ***taxonomically identical core microbiomes*** throughout the rest of the paper (**Fig. 1C**). The three taxa represent the most common species of the three most abundant bacterial genera in cheese rinds (17, 20). *Staphylococcus, Brevibacterium*, and *Brachybacterium* enter the dairy environment from the raw milk used for cheese production and therefore have the potential to co-occur and adapt to abiotic and biotic conditions within local cheese production facilities (21–23). Each species has a distinct colony morphology (**Fig. 1B**) making it easy to track composition in experimental communities. We predicted that intraspecific variation of core microbiome members across cheese rind communities would cause differences in community structure over time. We also predicted that strain-level diversity across core microbiomes would result in differences in community functions relevant for cheese aging, including pigmentation of the cheese rind biofilm and the production of aroma compounds.

**Figure 1:**
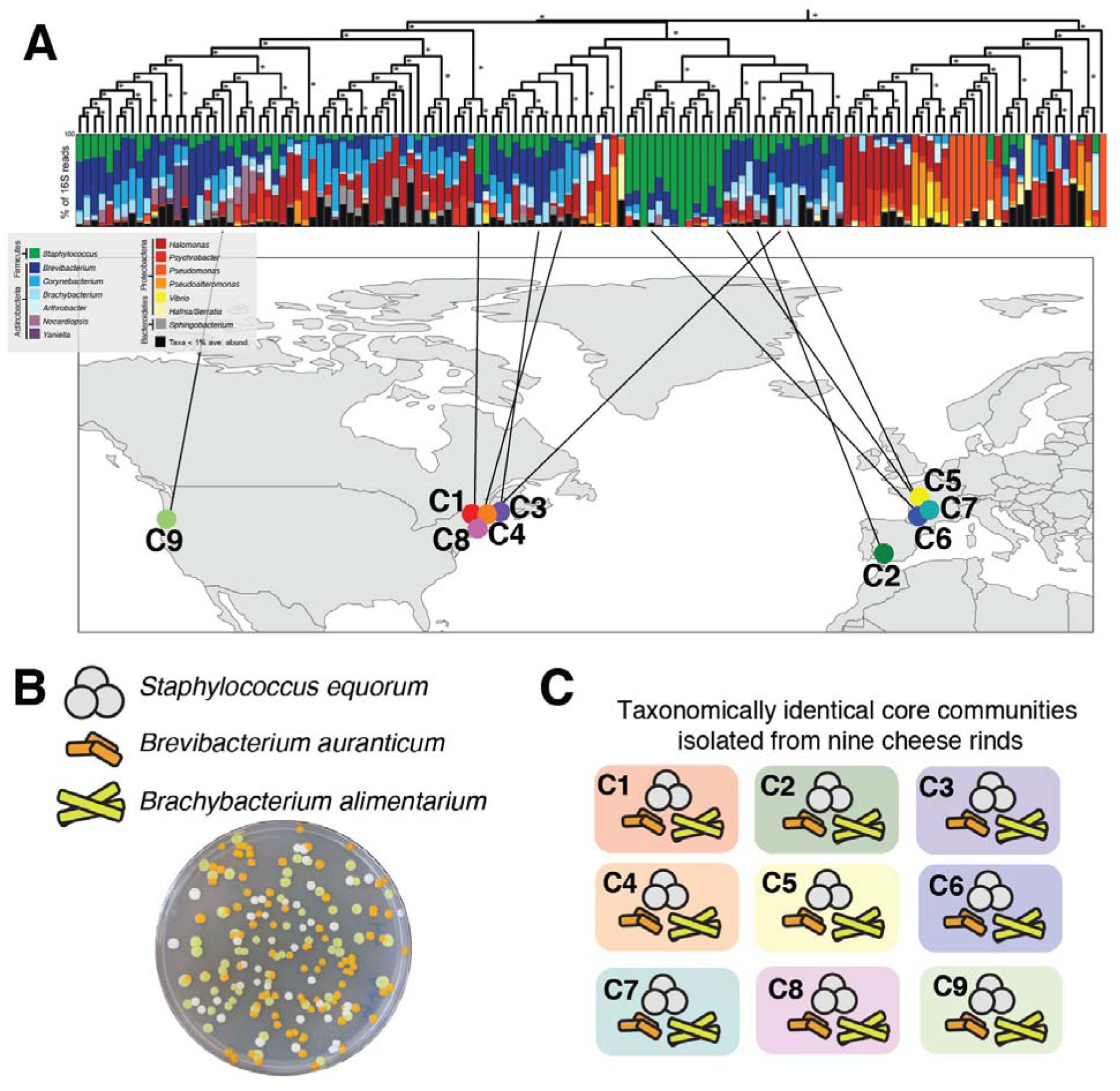
Isolation of nine taxonomically identical cheese rind core microbiomes. **(A)** The same three bacterial species – *Staphylococcus equorum*, *Brevibacterium auranticum*, and *Brachybacterium alimentarium -* were isolated from a set of 137 cheese rinds that were previously described using 16S rRNA gene amplicon sequencing (Wolfe et al. 2014). Each column represents average relative abundance data for one cheese rind microbiome. Data are clustered using an UPGMA tree based on Bray-Curtis dissimilarity. **(B)** The three core microbiome species have distinct colony morphologies. **(C)** Graphical representation of the nine core microbiomes as used throughout the manuscript.

## RESULTS

### Variation in genome content across taxonomically identical core microbiomes

To determine genomic variation across the nine taxonomically identical core microbiomes, we constructed draft genomes of each strain (**Table S1**). We used single-nucleotide polymorphisms (SNPs) in the core genes shared across all nine communities to determine phylogenomic divergence of each of the core communities (24). We then determined variation in functional gene content across the nine core communities using PGAP (25). For functional gene content analysis, we focused on accessory genes that were uniquely present in only one community as these genomic traits may help drive divergence in core microbiome functions.

Across the nine communities, 8,069 gene clusters were shared among all three species, making up the core metagenome of these communities. Using SNPs identified in this core metagenome with PanSeq, clear phylogenomic divergence across the nine cheese communities was apparent (**Fig. 2**). C1 was distant from the other eight core microbiomes, driven by the highly divergent *Staphylococcus* genome in this community. The eight other core microbiomes clustered into two broad phylogenomic groups: one containing C6 and C2, and the other containing the remaining six communities (**Fig. 2**).

**Figure 2:**
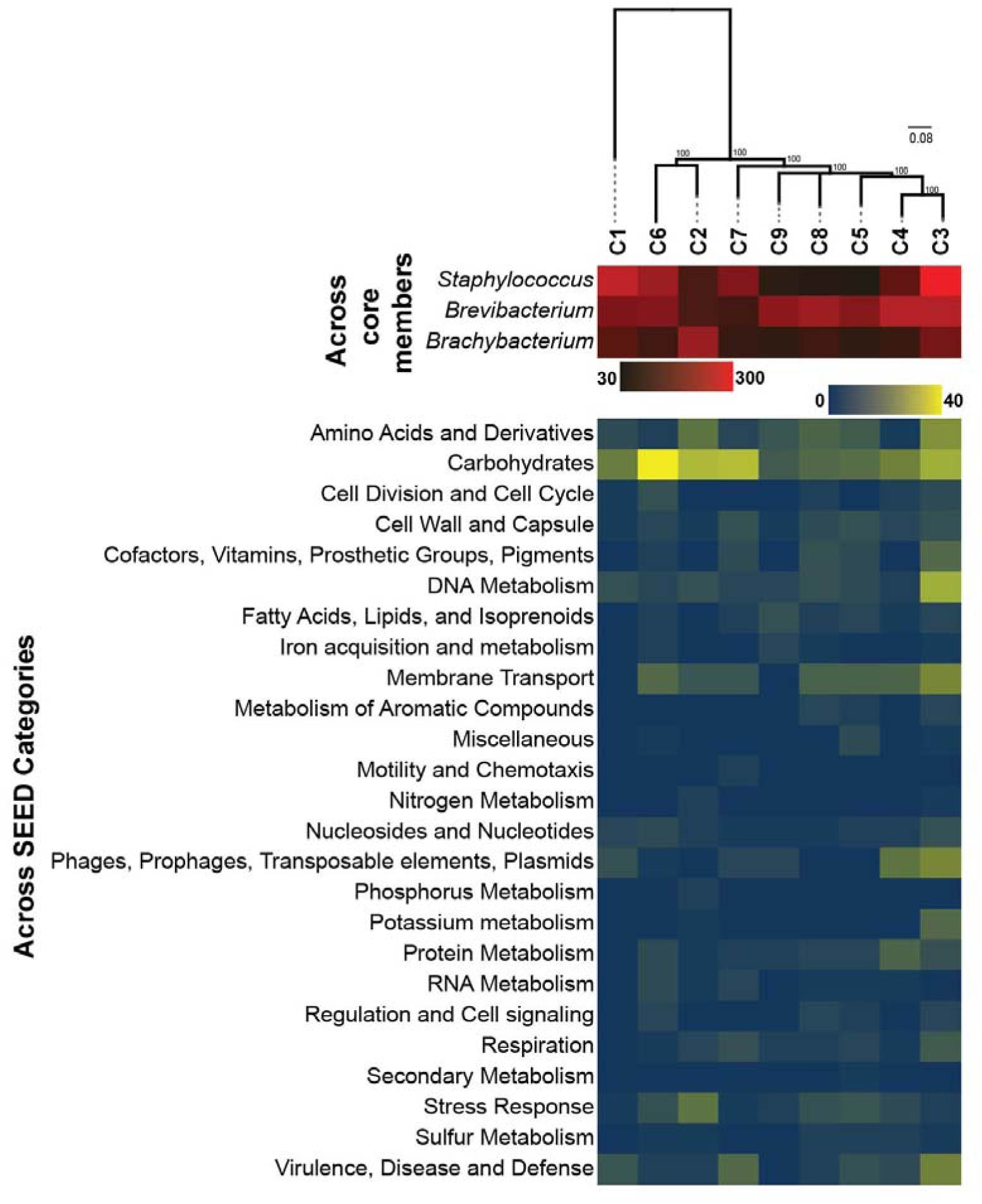
Accessory genome of the cheese rind core microbiomes. Heatmap indicates variation in the abundance of unique accessory gene clusters across the three individual taxa (top) and across SEED functional categories (bottom). Phylogeny is a maximum likelihood consensus tree constructed from SNPs identified across the nine core communities. Values are bootstrap support.

The total number of unique accessory gene clusters across the nine communities was highly variable, ranging from 246 (C5) to 630 genes (C3) (**Fig. 2**, **Table S2**). Variability in the abundance of accessory gene clusters was most prominent in *Staphylococcus* (ranging from 36-280 unique gene clusters across strains) and *Brevibacterium* (ranging from 72-213 unique gene clusters) suggesting that these taxa have the most dynamic accessory gene content in the cheese rind core metagenome.

Several biological processes were significantly enriched in core communities (**Table S3**). C3 had the most diverse enrichment of SEED categories, with overrepresentation of genes in potassium metabolism, carbohydrates, and DNA metabolism. Protein metabolism and phages/prophages/transposable elements/plasmids were overrepresented in C4. In C2, the accessory genome was significantly enriched with stress response genes. Carbohydrate-related genes were enriched in the C6 core microbiome. Some of these unique accessory genes could be functionally significant in the cheese rind environment. For example, *Brevibacterium* of C3 has a unique potassium transport system with high similarity to the *kdfABCF* operon (**Table S2**) that is known to play a role in salt stress in bacteria (26).

Collectively these genomic data demonstrate that taxonomically identical core microbiomes isolated from distinct cheeses are phylogenomically diverse and have variable genome content. Although the presence/absence of genes does not indicate actual functional potential of microbes, these comparative genomic data suggested to us that there could be divergence in how each taxa functioned within each community and how they responded to perturbations.

### Community assembly dynamics vary across taxonomically identical core microbiomes

We next determined whether strain-level differences impacted how the cheese rind communities assemble. A typical community succession in our lab model involves the following steps: 1) early colonization of *Staphylococcus* that can tolerate the low pH (5.0-5.2) of the cheese curd, 2) growth of *Brachybacterium* in middle succession, and 3) dominance by *Brevibacterium* at the end of succession (17, 27). We predicted two different potential impacts of strain-level variation on community assembly. In one scenario, distinct strains of *Staphylococcus*, *Brachybacterium*, and *Brevibacterium* across the nine communities may vary in genome content or growth rates in isolation, but these differences may be too minor to impact the dynamics of assembly of the three-member community. In this case, we expected nearly identical community composition across the different core microbiomes as strains of each species behaved similarly. Alternatively, strain-level differences may translate into differences in interactions with other community members or rates of growth within the community succession. In this scenario, we expected to observe reproducible changes in the composition of the communities as they assembled and differences in functional outputs.

To determine how strain-level differences across communities impact assembly dynamics, we used *in vitro* community assembly assays to measure total colony forming units (CFU) and community composition (relative abundance of each species) (**Fig. 3A**). Communities were quantified at three and ten days after inoculating equal amounts of each of the three bacterial species on the surface of cheese curd agar. Our previous work has demonstrated that this assay mimics *in situ* community dynamics (17, 27). We acknowledge that real cheese rind communities would develop over much longer time scales (weeks to months). In the context of this work, we used the community assembly assay in a standardized environment to demonstrate the *potential* for divergence in community assembly.

**Figure 3.**
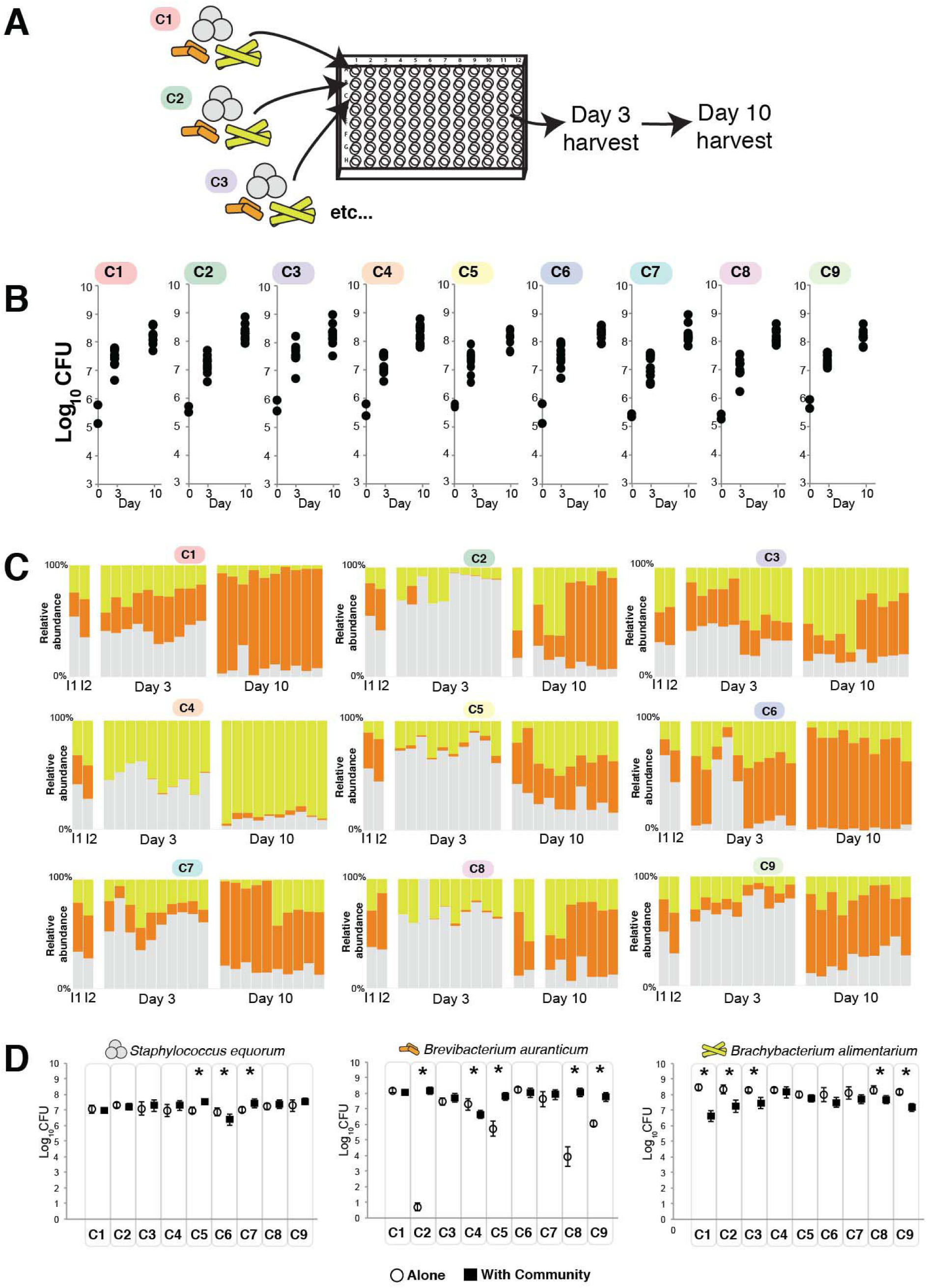
(previous page): Divergent community assembly across the nine cheese rind core microbiomes. **(A)** Experimental setup. Each set of three species from each core microbiome was inoculated into wells of 96-well plates. Communities were harvested three and ten days after inoculation. **(B)** Total community abundance as measured by CFUs of each of the nine core microbiomes. n=5 across two experimental replicates. **(C)** Relative abundance of each of the three bacterial species across each of the nine core microbiomes. Each column represents a replicate. I1 and I2 indicate the input compositions for the two independent experimental replicates. In the Day 3 and Day 10 datasets, the first five columns are from one experimental replicate and the second five are from a second experimental replicate. Blank columns represent replicates that were lost due to contamination. **(D)** Growth of each of the community members alone (open circles) and in the presence of the community (closed black squares). Each point represents the mean CFUs of the taxa and the error bars represent one standard deviation of the mean. Asterisks indicate significant differences between growth alone and growth in the community (n=5, t-test, *P* < 0.05).

At both three and ten days of community assembly, there were nearly no differences in total community abundance as measured by combined CFU of all three species (**Fig 3B**, Day 3 ANOVA *F*_8,81_ = 2.07, *P* = 0.05; Day 10 ANOVA *F*_8,79_ = 0.46, *P* = 0.88). However, there were substantial differences in community composition across the nine core communities (Day 3 permutational multivariate analysis of variance [PERMANOVA] *F* = 4.005, *P* = 0.0001; Day 10 PERMANOVA *F* = 5.57, *P* = 0.0001). Many communities (C1, C2, C6, C7) were dominated by *Brevibacterium* at the end of succession (**Fig 3C**). Some communities had a relatively even mix of all three species (C5, C3, C8, and C9). Community C4 had a very dissimilar structure with a high abundance of *Brachybacterium* at the end of succession and a low abundance of *Brevibacterium*.

A simple explanation for differences in community composition across the nine core communities is that individual bacterial strains have different growth abilities alone and in the community. Those taxa and strains that grow best alone and with the community present should be the most abundant members of the community. To test this, we determined total growth of each of the 27 strains on cheese curd agar and compared growth alone after ten days to growth in the community. All *Staphylococcus* species grew well alone and had limited responses to growth in the community (**Fig. 3D**). Two strains were slightly stimulated by growth in the community (C5 and C7) and one was slightly inhibited (C6). In contrast to the relatively even growth of the *Staphylococcus*, the *Brevibacterium* strains had variable growth alone across the nine core communities. Four of the *Brevibacterium* strains grew poorly by themselves on cheese curd agar (C2, C5, C8, and C9) and were strongly stimulated by growth in the community. One *Brevibacterium* strain (C4) was inhibited by growth in the community. All *Brachybacterium* strains grew well on cheese curd by themselves and were generally inhibited when grown in the community.

For all three taxa, mean growth alone was a very poor predictor of mean relative abundance in the community (*Staphylococcus* r^2^ = 0.166, *P*= 0.276; *Brevibacterium* r^2^ = 0.001, *P* = 0.923; *Brachybacterium* r^2^ = 0.020, *P* = 0.716). A somewhat better predictor of mean relative abundance was how growth of each strain was impacted by the community (*Staphylococcus* r^2^ = 0.672, *P*<0.01; *Brevibacterium* r^2^ = 0.013, *P* = 0.773; *Brachybacterium* r^2^ = 0.319, *P* = 0.113). This suggests that interactions between each of the strains and their communities may contribute to differences in community composition across the nine core microbiomes. For example, the inhibition of *Brevibacterium* and lack of inhibition of *Brachybacterium* in C4 may partly explain why this community was the only one to be dominated by *Brachybacterium*.

### Variation in responses to abiotic and biotic perturbation across core microbiomes

Core microbiomes may experience abiotic or biotic perturbations that could alter community assembly and function. We predicted that if individual core members have evolved different responses to stress or if the communities have coevolved stress-response mechanisms, taxonomically identical core microbiomes may have divergent responses to perturbations. Two major perturbations in cheese rind core microbiomes are salt and interactions with fungi (17, 20, 28). Salt concentrations are initially high on the surface of fresh cheese because salt is applied to the cheese surface or via a brine (29). The salt diffuses into the cheese and eventually equilibrates to around 3% salt in the rind environment of many cheeses. Core cheese rind microbiomes also experience interactions with fungi, ranging from yeasts (e.g. *Debaryomyces* and *Galactomyces* species) to molds (*Fusarium*, *Scopulariopsis*, and *Penicillium* species) (17, 20, 30). *Penicillium* species are widespread in cheese rinds and can strongly inhibit diverse cheese rind bacteria (17, 27, 31) potentially through the production of secondary metabolites or other mechanisms.

To determine how the nine core microbiomes respond to salt and fungal perturbations, we used the same community assembly assay described above with the addition of two treatments: a 6% NaCl treatment and a +*Penicillium* treatment. We used a strain of *Penicillium* that was isolated from a natural rind cheese and was previously demonstrated to inhibit cheese rind bacterial growth (17). Across isolates of all three taxa, both the 6% NaCl and +*Penicillium* treatments caused a general decrease in total growth across all nine core microbiomes with +*Penicillium* causing stronger growth inhibition (**Fig. 4A**). Core microbiomes had variable responses to the two perturbations. The *Penicillium* perturbation caused the most significant shifts in community composition with six out of nine core communities showing significant changes in community composition (**Fig. 4B-C**). In some communities, *Penicillium* caused a major increase in *Brachybacterium* relative abundance (C2 and C3). In others, *Penicillium* caused an increase in the relative abundance of *Staphylococcus* (C1, C8, and C9). The 6% salt treatment caused fewer shifts in community composition with only two communities (C5 and C6) responding to the higher salt environment. In both cases, *Brevibacterium* increased in relative abundance.

**Figure 4.**
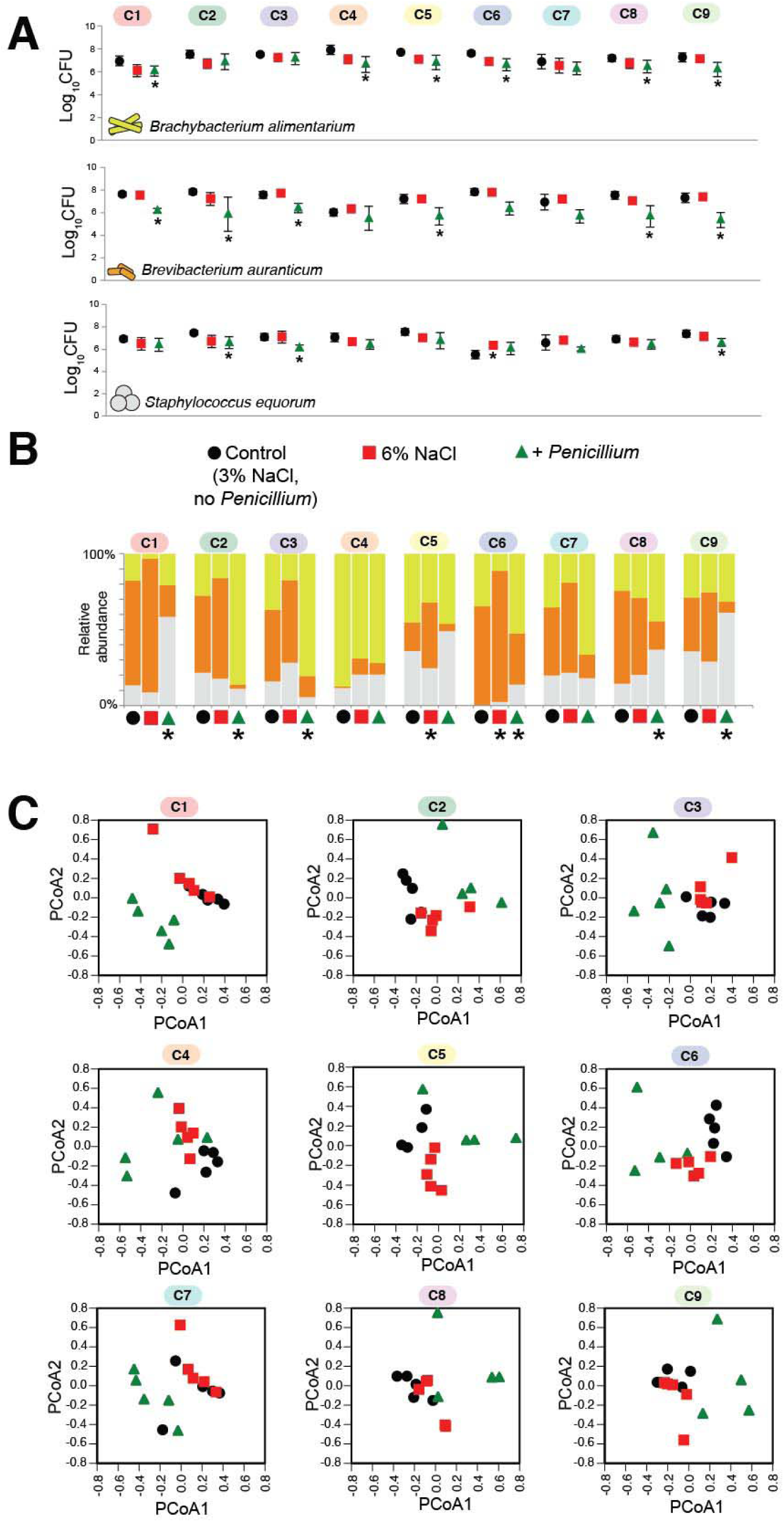
(next page): Response of the nine cheese rind core microbiomes to abiotic and biotic perturbations. **(A)** Responses of each taxa to abiotic (6% salt) and biotic (*Penicillium*) disturbance. Each point represents the mean CFUs of the taxa in that community at Day 10 (n=5) and the error bars represent one standard deviation of the mean. Asterisks indicate significant difference in growth compared to control based on Kruskal-Wallis test (p<0.05). (B) Mean community composition in the three treatments. Asterisk indicates significant difference in community composition compared to control based on PERMANOVA. (C) Principal coordinates analysis of replicate communities in the three treatments. PCoA is based on Bray-Curtis dissimilarity of absolute abundances of each community member.

### Strain-level diversity of cheese rind core microbiomes drives divergent pigment and aroma production

Our experiments above demonstrate that strain-level diversity of the core cheese rind taxa drives divergence in community composition across the nine core microbiomes. Does this divergence lead to cheeses with different properties that could be perceived by consumers? Differences in community composition may not necessarily translate into differences in functional outputs. Many studies of the microbiome have suggested that communities with different compositions may have similar functions due to functional redundancy across community members (32–34). While our comparative genomic analysis above suggested potential functional differences across the cheese communities, many of the core community functions were conserved in the core genome and variation in accessory genes may have little impact on community functions. To determine whether divergence in composition of the core microbiomes also translated into differences in functional outputs, we measured two important traits of cheese rind microbiomes: rind color and volatile organic compound (VOC) production.

Cheese rind bacteria define how the cheese appears to customers through the production of cellular pigments such as carotenoids or the secretion of pigmented extracellular metabolites into the curd (35–39). The three bacteria in our model community produce distinct pigments (**Fig. 1B**) and shifts in their relative abundance could translate into changes in rind color. Using a colorimeter, we measured rind color after 10 days. Communities had significantly different color development (ANOVA F_9,39_ = 524.9, *P* <0.0001), with C3, C4, C6, C7, and C9 having significantly greater a* values compared to the control, indicating more red pigmentation (**Fig. 5A**). All communities had significantly greater b* values compared to the control (ANOVA F_9,39_ = 139.6, *P* <0.0001), with C3 and C4 having the greatest values and appearing the most orange (**Fig. 5A**).

**Figure 5:**
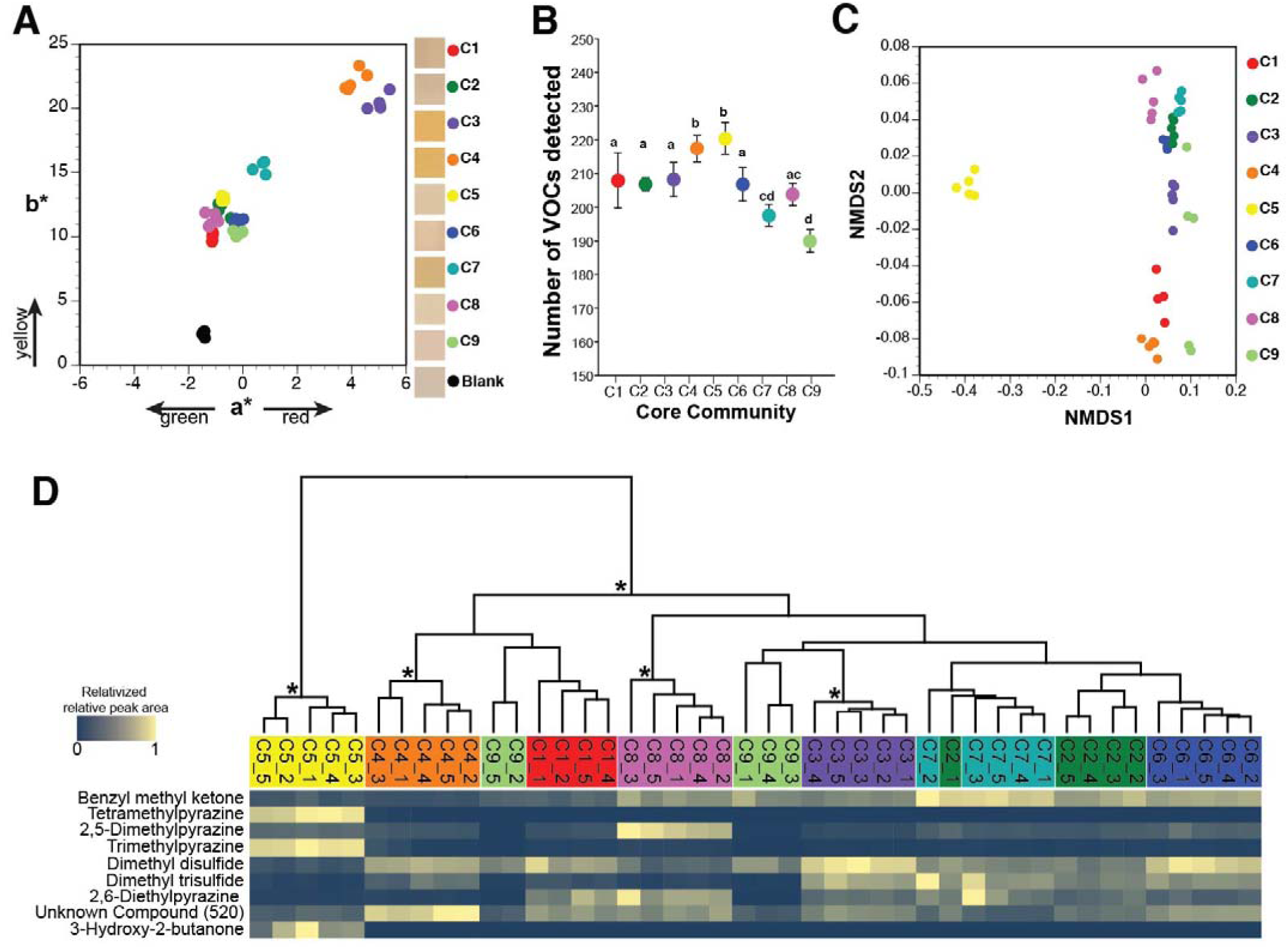
Functional diversity across nine cheese rind core microbiomes. **(A)** Color profiles of experimental rind communities after ten days of rind development. Each dot represents a replicate cheese rind community (n=5). Boxes in legend are representative photos of the experimental cheese surface from each community. **(B)** Total volatile organic compound (VOC) diversity across the nine cheese communities. Each point represents the mean number of VOCs detected in each community and the error bars represent one standard deviation of the mean (n=5). Core communities that share the same letter are not significantly different from one another based on Kruskal-Wallis test (p<0.05)**. (C)** Non-metric multidimensional scaling of total VOC profiles. Each dot represents a replicate cheese rind community (n=5). **(D)** Relative abundance of VOCs that contributed the most to the Bray-Curtis dissimilarity across communities (as determined by SIMPER analysis). Because total concentrations of VOCs are highly variable across different compounds, visualization was simplified by relativizing the relative peak area from GC-MS chromatograms within each VOC to the highest concentration detected for that VOC. Data are clustered together by total VOC profiles using a UPGMA tree. Asterisks indicate clusters with > 70% bootstrap support.

As the rind biofilm decomposes fats, proteins, and other components of the cheese substrate, a diversity of VOCs are produced that are aromatic (40–42). Using headspace sorptive extraction (HSSE) followed by gas chromatography-mass spectrometry (GC-MS) analysis (43, 44), we quantified VOCs produced by each community after 10 days of cheese rind development. Across all nine communities 248 unique VOCs were detected with significant differences in the mean VOCs per community (**Fig. 5B**, ANOVA F_8,35_ = 28.9, *P* <0.0001). The composition of VOCs across the nine cheese communities was significantly different (**Fig. 5C**, PERMANOVA *F* = 62.38, *P* < 0.001, **Table S4**). Using a SIMPER analysis, nine compounds contributed more than 1% to the average overall Bray-Curtis dissimilarity: benzyl methyl ketone (27% contribution; odor = floral/fruity), tetramethylpyrazine (19%; odor = nutty/musty/chocolate/coffee), 2,5-dimethylpyrazine (13%; odor = nutty/musty/chocolate/coffee), trimethylpyrazine (12%; odor = nutty/musty/chocolate/coffee), dimethyl disulfide (9%; odor = sulfurous/cabbage/onion), dimethyl trisulfide (2%; odor = sulfurous/cabbage/onion), 2,6-diethylpyrazine (2%; odor = nutty/musty/chocolate/coffee), unknown compound 520 (1%; odor = unknown), and 3-hydroxy-2-butanone (1%; odor = sweet/buttery/creamy). C5 had the most distinct VOC profile of all communities with high amounts of tetramethylpyrazine, trimethylpyrazine, and 3-hydroxy-2-butanone and low amounts of the major sulfur compounds, suggesting a nuttier and more buttery aroma profile.

## DISCUSSION

Using taxonomically identical three-member communities isolated from nine distinct cheeses, our work demonstrates the significance of strain-level variation for microbiome community assembly and function. Studies of plant and animal communities have demonstrated that intraspecific genetic and phenotypic diversity can impact community assembly and function (45–47). Here we demonstrate that intraspecific diversity of taxonomically identical core microbiome members can impact the relative abundance of community members as well as functional outputs of the communities. Many communities did converge on a similar composition despite having substantial variation in accessory gene content. But several communities had substantially different structures and functions even though the initial inoculum was identical. Some communities had relatively even coexistence of the three community members, while others were dominated by either *Brevibacterium* or *Brachybacterium*. The divergence was not due to stochastic community assembly across replicates as we observed highly reproducible community structures across replicate experiments.

The goal of this work was to determine whether taxonomically identical core microbiomes have similar community dynamics and functions. The limited number of core communities (nine) makes it difficult to pinpoint specific ecological or genetic mechanisms that may be underlying the observed differences across the core communities. One simple explanation for the dominance of different taxa across the core microbiomes is differences in growth of individual strains. Our experiments comparing growth alone versus in the community demonstrates variable growth rates and interactions with the community for each of the three taxa. However, it does not fully explain community structure. For example, in C4 where *Brachybacterium* dominated, the *Brachybacterium* strain had similar levels of growth alone and interactions with community members as other communities where *Brevibacterium* dominated (e.g. C5 and C6). Future work exploring the roles of inhibitory and cooperative interactions will pinpoint specific mechanisms explaining the variable community assembly dynamics of cheese rind core microbiomes.

The evolutionary processes that have generated the divergent species and community-level responses of our core cheese microbiomes are currently unknown. It is possible that each core microbiome has experienced different evolutionary histories in each cheese production environment. As new batches of cheese are introduced to a cave environment, communities may be repeatedly transferred to these new cheeses. This repeated colonization of the cheese substrate could allow each of the core microbiomes to evolve collectively as a community in the individual production environments (48). Each environment may have unique abiotic selection pressures, including salt concentrations, milk composition, and temperature that could shape the evolutionary trajectories of these communities. The core microbiomes could also experience highly divergent biotic environments. For example, these core communities were isolated from cheeses with variable fungal environments, ranging from yeast to filamentous fungi (17). Future work using experimental evolution to attempt to create divergent communities from an ancestral core microbiome should begin to help us understand the drivers of core microbiome diversification.

Our model communities represent the widespread bacterial taxa found in cheese rinds. We acknowledge that these communities have several constraints that may impact translation of our results to other systems. First, our communities only had three bacterial species. While some widespread microbiomes have low species diversity (5, 49), many microbiomes have much higher levels of diversity. Would taxonomically identical core microbiomes with higher taxonomic diversity also demonstrate divergence in assembly and functions? With greater potential for higher-order interactions and a higher number of potential functions with increasing species diversity, we predict that increasing diversity may lead to even more divergent communities. Our model communities also used a single strain of each species within each core microbiome. In constructing our communities, we chose to ignore potential intraspecific variation within each of the nine core communities and assumed that the isolated taxa represented the most common genomic type of the species within each of the core communities. Metagenomic sequencing studies have identified multiple co-existing strains of the same microbial species (3, 16, 56–60) and these strains may interact with each other and other community members to impact community composition. It would be fascinating to see how including intraspecific diversity within core microbiomes may impact community assembly and function.

In a large amplicon-sequencing study of cheese rind microbiomes, we demonstrated that taxonomically identical cheese rind communities could form in very different cheese-making regions (17). This was surprising given that these cheeses have divergent sensory properties. Many of these differences could be driven by ingredients, length of aging, or other cheese processes. Our current findings suggest that the variability in the qualities of surface-ripened cheeses could also be driven by strain-level differences across the cheese communities. We acknowledge that our lab cheese rinds are not real cheeses and only represent potential patterns of cheese rind community assembly. But it is very likely that the differences observed across the nine core microbiomes would translate to actual cheese production. Previous studies of fermented food microbes have pointed out strain-level differences of individual species used in fermented foods (50–54), but studies demonstrating the functional significance of strain-level variation at the community level are rare (55). To help preserve the unique identities of cheeses made in specific regions, it may be helpful for cheese producers to identify the unique genomic and functional properties of their core microbiomes and maintain these communities.

More broadly, our work in these model microbiomes may have implications for both the design and management of core microbiomes in other systems. First, our work demonstrates that taxonomic profiling of microbiomes may not provide useful predictors of assembly dynamics and functions. Amplicon based approaches of sequencing microbiomes, such as using 16S rRNA gene sequencing, only capture high-level taxonomic diversity. As we have demonstrated, taxonomically similar communities can have very different dynamics. Fortunately, microbiome sequencing studies are moving toward shotgun-metagenomic approaches that could capture the strain-level diversity that we observed across our nine communities (3, 16, 56–60). Our work also suggests that it might be hard to predict microbiome responses to disturbances using taxonomic profiles alone. For example, across individuals that have similar skin core microbiomes, responses to environmental stresses such as antibiotics may depend on the specific strains and genomic content of the core communities. Finally, when designing synthetic microbiomes, our work suggests that the individual ‘parts’ (strains of species) may alter desired outcomes.

## METHODS

### Isolation and maintenance of core microbiome members

Frozen glycerol stocks of communities initially characterized using metagenomic sequencing (Wolfe et al. 2014) were plated out on plate count agar with milk and salt (PCAMS) to culture bacteria. Colonies with morphotypes that had the appearance of one of the three target species were streaked from single colonies. *Staphylococcus equorum* colonies are usually fast-growing, smooth, medium-sized, flat, and either white or light golden in color. *Brevibacterium auranticum* colonies are usually slow-growing, medium-sized, and orange. *Brachybacterium alimentarium* colonies have medium growth rates, are large and flat, and are yellow-green in color. Initial identification of the isolates was done using the 16S rRNA region using primers 27f and 1492r.

### Comparative genomics

The genome of each bacterial strain was sequenced, assembled, and annotated as we previously described for *Staphylococcus* species (27). Briefly, DNA was extracted using MoBio PowerSoil DNA extraction kits from pure cultures grown for one week on PCAMS. Approximately 1 µg of purified gDNA was sheared using a Covaris S220 to approximately 450 base pair lengths and was used as the input for a New England Biolabs NEBNext Ultra DNA Library Prep Kit for Illumina. Libraries were spread across multiple sequencing lanes with other projects and were sequenced using 100 base-pair, paired-end reads on an Illumina HiSeq 2500. Approximately 10 million reads were sequenced for each genome. Failed reads were removed from libraries and reads were trimmed to remove low quality bases and were assembled to create draft genomes using the *de novo* assembler in CLC Genomics Workbench 8.0. Assembled genomes were annotated using RAST(61). All genome assemblies have been deposited in NCBI (accession numbers in Table S1).

To identify phylogenomic relationships between each of the nine core communities, we used PanSeq (24) to identify SNPs across the core genome of each of the nine genomes for each of the three species. A SNP file for each species from each community was then concatenated together to create a community SNP file. RAxML 8.2.11 (with GTR GAMMA nucleotide model and 100 bootstrap replicates) was used to create a maximum likelihood phylogeny of the nine communities using the SNP file.

To compare the presence and absence of genes across strains and species, core and accessory genes were identified by assigning protein-coding sequences to functionally orthologous groups using the MultiParanoid method of the PanGenome Analysis Pipeline (PGAP) (25). Species-to-species orthologs were identified by pairwise strain comparison using BLAST with PGAP defaults: a minimum local coverage of 25% of the longer group and a global match of no less than 50% of the longer group, a minimum score value of 50, and a maximum E value of 1E−8. Multistrain orthologs were then found using MultiParanoid (80). Enrichment of SEED subsystem categories in each of the nine core communities was determined using Fisher’s exact test with false-discovery rate correction.

### Community assembly assays

To measure assembly of the distinct core communities, approximately 20,000 CFU of each species was inoculated on the surface of 150 µL of cheese curd agar (3% salt) distributed into replicate wells of a 96-well plate, as previously described (17, 27). Communities were incubated aerobically at 24°C in the dark, and harvested at 3 and 10 days after inoculation, which represent early and late community succession (17). To determine community composition of individual replicate communities, the community was pestled in 600 µL of 1X phosphate buffered saline, serially diluted, and plated onto PCAMS. PCAMS plates were incubated for a week before counting the abundance of each bacterial species. To measure growth alone, the same density of CFU of each taxa alone was inoculated into wells. Five technical replicates of each community were performed in each of two experimental replicates.

Salt (6%) and fungal (+*Penicillium*) perturbation experiments were conducted using the same community assembly assay, but with 6% salt cheese curd agar or with the addition of *Penicillium*. *Penicillium* strain #12, isolated from a natural rind cheese in Vermont, was used in these experiments. We used this strain because it was isolated from a cheese where the *Staphylococcus*, *Brachybacterium*, and *Brevibacterium* were also found and it was used in previous experiments in our lab (27, 31). The exact species identification of this mold is unknown, but it belongs to section *Fasciculata* with other cheese *Penicillium* species. *Penicillium* was inoculated at an initial density of 2000 CFUs. Community composition in these experiments was determined as described above except that cycloheximide was added to PCAMS plates used for bacterial community isolation to eliminate fungal growth.

### Color and VOC analyses

To measure rind color and VOC production, we constructed larger versions of each of the nine core communities on cheese curd agar poured into Petri dishes (60mm wide) to allow for a larger sampling area. To construct the rind communities, 600,000 CFU of each species was inoculated across the surface of the cheese curd agar. Experimental cheeses were incubated for 10 days in the dark at 24°C before color and VOC analyses.

To measure differences in color of the experimental cheeses, we used a CTI A6CTI10 spectrocolorimeter. This handheld colorimeter uses the CIELAB color space to quantify both lightness (L*) and two chromatic coordinates (a* and b*). Similar colorimeters have been used to quantify cheese rind color (62). Higher values of a* (a*+) indicate red colors while lower values (a*-) indicate green colors. Higher values of b*(b*+) indicate yellow while lower values (b*-) indicate blue colors. Colorimeter readings were taken by placing a 30mm Petri dish lid upside down on the middle of the surface of the rind and then placing the colorimeter on the Petri dish surface. This was done to protect the colorimeter from the sticky rind surface and to avoid cross-contamination across replicates.

Cheese volatiles were collected from experimental cheese rinds by headspace sorptive extraction (HSSE) using a polydimethylsiloxane (PDMS) coated magnetic stir-bar. HSSE is an equilibrium-driven, enrichment technique in which 10mm long × 0.5 mm thick stir-bars, Twister^TM^ (Gerstel), were suspended 1 cm above the sample by placing a magnet on the top side of the collection vessel cover. Five replicates of each culture were sampled for four hours. After collection, the stir-bar was removed and spiked with 10 ppm ethylbenzene-d_10_, an internal standard obtained from RESTEK. The internal standard was used to determine the relative concentration of each compound. Organics were introduced into the gas chromatograph/mass spectrometer (GC/MS) by thermal desorption. In addition to Twister blanks, analysis of the cheese curd agar media was made to assess background interferences. Compounds present at equal or higher relative concentrations in the media compared to the samples were removed from the data.

Analyses were performed using an Agilent 7890A/5975C GC/MS equipped with an automated multi-purpose sampler (Gerstel). The thermal desorption unit (TDU, Gerstel) provided splitless transfer of the sample from the stir bar into a programmable temperature vaporization inlet (CIS, Gerstel). The TDU was heated from 40°C (0.70 min) to 275°C (3 min) at 600°C/min under 50ml/min of helium. After 0.1 min the CIS, operating in solvent vent mode, was heated from −100°C to 275°C (5 min) at 12°C/s. The GC column (30 m × 250 µm × 0.25 µm HP5-MS, Agilent) was heated from 40°C (1 min) to 280 °C at 5°C/min with 1.2 mL/min of constant helium flow. The MS was scanned from 40 to 350 *m/z*, with the EI source at 70 eV. A standard mixture of C7 to C30 n-alkanes (Sigma–Aldrich) was used to calculate the retention index (RI) of each compound in the sample.

The Ion Analytics spectral deconvolution software (Gerstel) was used to analyze the GC/MS data (63, 64). A target/nontarget data analysis approach was employed where previously constructed databases are used to identify target compounds in the sample based on spectra deconvolution of their irons and abundances. Once found, each compound’s mass spectrum was subtracted from the peak’s total ion current (TIC) signal. Each resulting peak scan was inspected to determine if residual ion signals were constant (±20%) or approximated background noise. If constant, the software recorded the retention time, mass spectrum, 3-5 target ions and their relative abundances into the database. Finally, sample data were compared to reference compound data in the database, viz., RI and MS (positive identification), or to commercial libraries and literature (tentative identification). Once assigned, the database was annotated to include compound name, CAS#, and RI. If neither positive nor tentative identification was possible (an unknown), a numerical identifier was used to identify the compound. The database was annotated to include the same GC/MS information described above. In contrast, if peak scans differed (an unresolved peak), the software searched for 3-5 invariant scans, averaged their spectra, and then subtracted the average spectrum from the TIC signal. This process was repeated until the residual signal at each scan approximated background noise. If peak signals failed to meet the user-defined criterion below, no additional information was obtained.

### Statistical Analyses

To determine differences in community composition with all core microbiome experiments, PERMANOVAs with Bray-Curtis dissimilarity were used. ANOVA on log-transformed data was used to determine significant differences between total CFU across experiments. In the cases of unequal variances (the individual taxa growth in perturbations), Kruskal-Wallis tests were used. To determine relationships between relative abundance and growth of individual strains, linear regressions were used. To compare total growth alone to growth in the community, t-tests were used. Differences in a* and b* values in the pigmentation assay were determined using ANOVA. To determine differences in VOC composition across the nine communities, PERMANOVAs on Bray-Curtis dissimilarity of relative peak area were used. A SIMPER analysis of relative peak area of VOCs was used to identify the contributions of each VOC to Bray-Curtis dissimilarity.

## ACKNOWLEDGEMENTS

Esther Miller, Grace Cox, Elizabeth Landis, Freddy Lee, and Megan Biango-Daniels provided very helpful feedback on earlier drafts of this manuscript. This work was supported by NSF grant 1715553 to B.E.W.

## AUTHOR CONTRIBUTIONS

B.E.W. isolated the bacterial strains from the nine original cheeses. B.A.N. designed, conducted, and analyzed all *in vitro* cheese experiments. N.K. and A.R. designed and performed the VOC data collection and analysis. E.K.K. and B.E.W. performed bioinformatic analyses. B.A.N. and B.E.W. performed statistical analyses on the community assembly and functional assays. B.A.N., E.E.K., N.K., A.R., and B.E.W. wrote the manuscript.

## SUPPLEMENTARY TABLES

**Table S1:** Overview of bacterial strains and genomes used in this study

**Table S2:** Distribution of gene clusters in the three taxa from each of the nine core microbiomes. When a cell is filled, it indicates that a predicted gene belongs to a gene cluster (row). In some communities, multiple genes belong to a single gene cluster. The identifiers in the cells are the gene IDs of each of the genomes based on the RAST annotation of that genome.

**Table S3:** Enrichment of SEED subsystem categories in core microbiomes based on Fisher’s exact test.

**Table S4:** Relative peak area of each volatile organic compound detected from the experimental cheese communities. The “_1, _2, etc.” indicates replicates within each of the nine core communities.

